# The Activity of the Stress Modulated Arabidopsis Ubiquitin Ligases PUB46 and PUB48 is Partially Redundant

**DOI:** 10.1101/2021.10.10.463813

**Authors:** Gal Zizelski-Valenci, Dina Raveh, Dudy Bar-Zvi

## Abstract

The Arabidopsis ubiquitin ligases PUB46, PUB47 and PUB48 are encoded by paralogus genes; *pub46* and *pub48* mutants display increased drought sensitivity compared to wild type (WT). Although the phenotype displayed in the single gene mutants, suggest that each has specific biological activity, PUB46 and PUB48 activity may be also redundant. To test functional redundancy between two gene products requires a double mutant. Unfortunately, the close proximity of the *PUB46* and *PUB48* gene loci precludes obtaining a double mutant by crossing the available single mutants. We thus applied microRNA technology to reduce the activity of all three gene products of the *PUB46-48* subfamily by constructing an artificial microRNA (aMIR) targeted to this subfamily. Expressing aMIR46-48 in WT plants resulted in increased drought-sensitivity, a phenotype resembling that of the single *pub46* and *pub48* mutants, and enhanced sensitivity to methyl viologen, similar to that observed for the *pub46* mutant. Furthermore, the WT plants expressing aMIR46-48 also revealed reduced inhibition by ABA at seed germination, a phenotype not evident in the single mutants. Expressing aMIR46-48 in *pub46* and *pub48* mutants further enhanced the drought sensitivity of each parental single mutant and of WT expressing aMIR46-48. Thus, whereas the gene-specific activity of the PUB46 and PUB48 E3s is partially redundant in that absence of either E3 leads to drought sensitivity, our ability to eliminate the activity of both *PUB46* and *PUB48* in the same plant reveals additional gene specific facets of their activity in the reaction to abiotic stress.

## Introduction

Plants being sessile lack the option of relocating to escape harsh conditions and have evolved physiological, developmental, biochemical, and molecular mechanisms to endure periods of stress. Plant exposure to abiotic stress evokes a global shift in gene expression to achieve a new proteostasis in which both protein translation and degradation are specifically modulated. The Ubiquitin Proteasome System (UPS) is a central, highly regulated mechanism for protein degradation in all eukaryotes ^1,2^. Proteins are ubiquitylated in a three-step cascade comprising E1 ubiquitin activating, E2 ubiquitin conjugating and E3 ubiquitin ligase enzymes. The ubiquitylated proteins are bound by the 26S proteasome; the ubiquitin tag is removed and recycled, and the target protein is denatured and transferred to the 20S catalytic chamber of the proteasome for degradation into short peptides. The E3s recruit specific substrates and bind the E2 and transfer UB molecules onto lysine residues of the substrate. Thus they determine the half-life of particular protein substrates. Plant genomes contain an exceptionally high number of genes encoding components of the UPS: over 1,600 UPS loci, almost 6% of the Arabidopsis genome encode UPS proteins several times that of other organisms from yeast to human ^1^. The E3s Ub recognize the degradation substrates conferring specificity and over 1,400 Arabidopsis genes encode putative E3s suggesting that this high number is essential for maintaining functional plants.

E3s can be grouped into a number of subclasses based on their motifs and oligomeric forms ^3^. One of the smaller gene families comprises Plant U-box (PUB) E3s defined by a highly conserved ∼70 amino acid long U-box domain first identified in yeast Ufd2 ^4^. The U-box fold is similar to the E3 RING domain ^5^ but is stabilized by salt-bridges rather than zinc-binding motifs ^6^. Unlike yeast, which has two U-box proteins, plants have a large number of genes encoding these putative E3: the Arabidopsis genome encodes 64 PUB ^7,8^ in addition to ∼700 putative F-box and 470 RING E3s ^9,10^. Many plant genomes have been analyzed for PUBs; gene numbers range from 56 in grapevine to 213 in wheat ^7,11-21^. In contrast, only 8 U-box E3 genes are present in the human genome ^22^. PUBs are divided into subclasses based on their domain composition ^23,24^. The largest class of Arabidopsis and rice PUBs contains ARM-repeat protein-protein interaction domains ^25-27^. U-box-ARM proteins are unique to plants ^28^ and PUB-ARM E3s play a role in diverse biological processes, including development, immunity and abiotic stress (reviewed by ^8^).

We have studied the paralogous Arabidopsis *PUB46, PUB47* and *PUB48* genes ^29,30^, particularly *PUB46* and *PUB48* as they are more highly expressed and their mutants give a strong hypersensitivity to water stress. The three genes are located in tandem on chromosome 5 and probably resulted from a recent gene duplication ^29^. Many plant genomes have undergone multiplication resulting in high numbers of gene families encoding homologous proteins ^31,32^. Surprisingly, despite encoding highly homologous proteins, a single T-DNA insertion mutant of either *PUB46* or *PUB48* results in hypersensitivity to water stress, indicating that each of these genes is essential for the drought response in a gene specific manner ^29^. However, the biological roles of *PUB46* and *PUB48* may also be partially redundant and these two E3s may share some substrates.

Gene multiplication often results in functional redundancy and to obtain a mutant phenotype in redundant genes it is necessary to produce double or multiple gene mutants. The physical proximity of the paralogous *PUB46, PUB47 and PUB48* genes precludes obtaining these by crossing the single mutant plants. However, due to the high homology of these genes we were able to employ RNA interference technology ^33^ to severely reduce the activity of all three *PUB46-48* genes in a single plant and to test whether *PUB46* and *PUB48* function is redundant in the response to abiotic stress. In RNAi, double stranded (ds) RNA is digested by a DICER-family RNase to yield 21-24 bp RNA products. One strand is then bound by an AGRONOUT (AGO) protein that is subsequently bound by the RNA-Induced Silencing Complex (RISC). In plants RISC associates with the target mRNA and either degrades it or inhibits its translation (reviewed by ^33,34^. RNAi comprises two classes of small RNAs: siRNA, small interfering RNAs that have perfectly complementary sequences and are prevalent in animals, and microRNA (miRNA), an imperfect dsRNA resulting from a genome-encoded short highly structured non-coding RNA precursor ^34^. Schwab et al developed a method for constructing artificial miRNA (aMIR) by introducing the desired siRNA sequence into the backbone of a natural plant miRNA. This allows design of an aMIR that can be targeted to a single mRNA or to a group of homologous mRNAs of choice ^35^.

Here we report that expression of a single artificial microRNA (aMIR46-48) that targeted all three *PUB46, PUB47* and *PUB48* genes in wild type (WT) plants led to uncreased drought sensitivity. Expressing aMIR46-48 in *pub46* and *pub48* mutants enhanced their drought sensitivity which exceeded that of the WT-aMIR46-48 plants. The creation of artificial triple *pub46-48* mutants exposed facets of their role in the abiotic stress response that are not redundant. Overexpressing aMIR46-48 also resulted in a reduced inhibition of germination by ABA, a phenotype not observed in the single mutants. Germination of WT expressing aMIR46-48 was also less sensitive to methyl viologen (MV). Thus, the creation of artificial triple *pub46-48* mutants exposed facets of their role in the abiotic stress response that are both redundant and nonredundant.

## Results

### Design of aMIR to silence *PUB46, PUB47* and *PUB48*

The high sequence homology of the *PUB46*-*PUB48* gene cluster allowed us to design a single aMIR sequence termed aMIR46-48. This aMIR46-48 targets the mRNA sequences encoding amino acids close to the N-terminus of each protein (nucleotides 140-160 of *PUB46* and *PUB48* and nts 119-139 of *PUB47*), and is predicted to be a high potency candidate sequence for aMIR affecting all three genes with a hybridization energy ranging between − 31.56 and −38.76 kcal/mol which should achieve stable hybridization with the target mRNAs. In addition, aMIR46-48 is not expected to have any off-targets. The designed sequence was included in the backbone of MIR319A ^35^ (Fig.1 and Fig. 1S) and then cloned into the plant transformation vector pCAMBIA 99-1 downstream of the constitutive 35S Cauliflower Mosaic Virus (35S CaMV) promoter. The resulting plasmid was used to transform Arabidopsis plants.

**Fig. 1.**
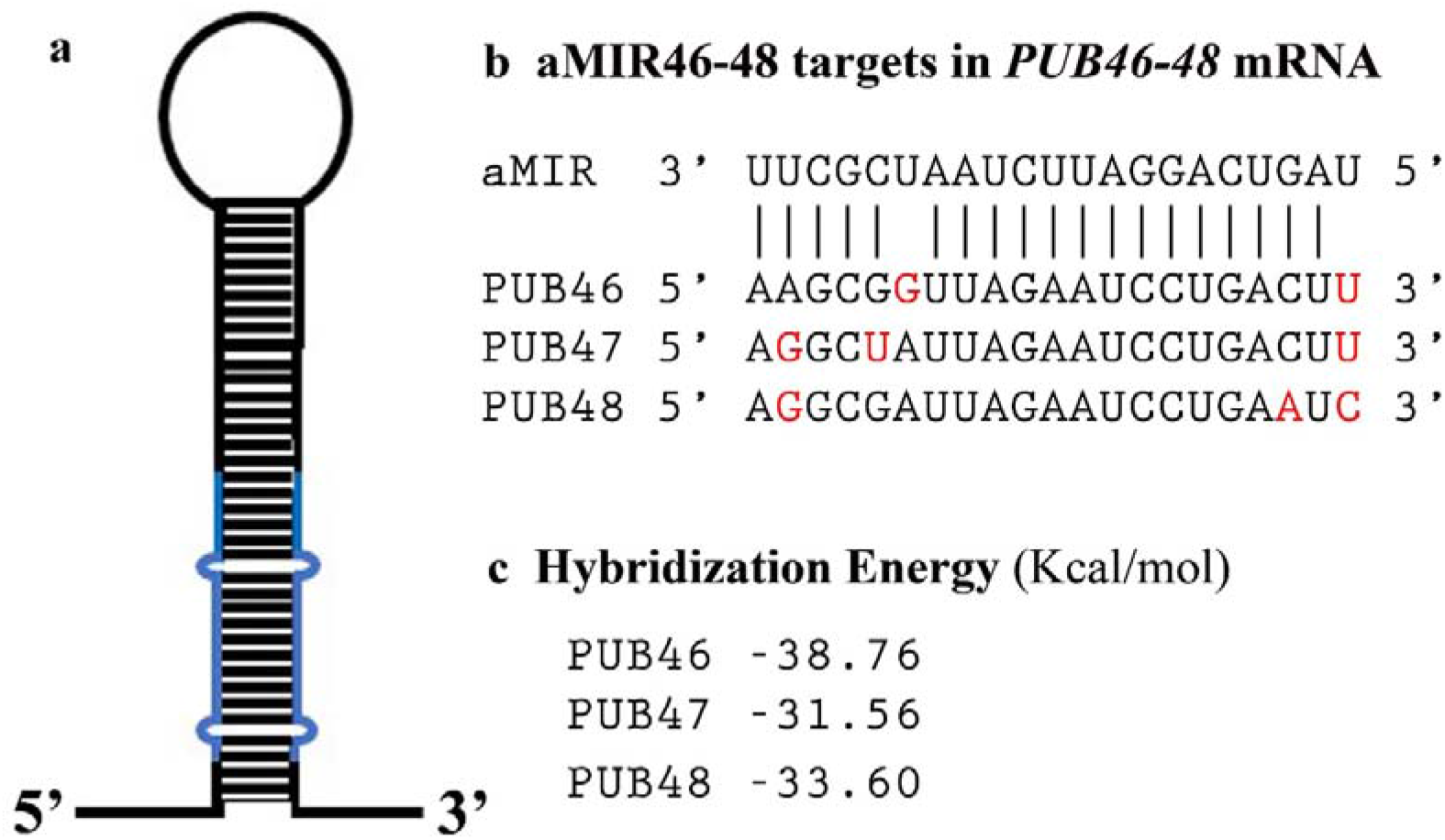
Diagram of aMIR46-48 and its targets in *PUB46-48* mRNAs. a. Diagram of aMIR46-48. Sequences in blue represent the two strands of sequence designed to specifically target the mRNA encoded by *PUB46, PUB47* and *PUB48*. b. Proposed hybridization by aMIR46-48 and the respective mRNA target sequences. Mismatches between aMIR46-48 and each target sequence are shown in red. The top two sequence lines show the hybridization between aMIR46-48 and *PUB46* mRNA. The respective sequences in *PUB47* and *PUB48* are aligned in the bottom two lines. c. Hybridization energy calculated for aMIR46-48 and its target mRNAs.

### RNA interference of *PUB46 PUB47* and *PUB48* in WT plants

The observed drought hypersensitivity of the *pub46* and *pub48* single gene mutants ^29^ serves as an unequivocal assay for effectiveness of gene silencing by aMIR46-48 in WT plants. We focused on *PUB46* and *PUB48* in our experiments as *PUB47* is expressed at a considerably lower level and *pub47* mutants do not show a stress related phenotype ^29^. We transformed WT plants with aMIR-46-48 and assayed four homozygous lines from independent transformation events each with a single T-DNA insertion. The phenotype of the aMIR46-48 transformants resembled WT plants when grown under optimal conditions on agar plates with 0.5 X MS medium or in pots. However, when rosettes of pot-grown irrigated plants were exposed to drought stress by withholding irrigation the plants expressing aMIR46-48 were hypersensitive to drought compared to the WT plants (Fig. 2). The observed drought-hypersensitivity of WT expressing aMIR46-48 resembled that of the *pub46* and *pub48* single genes ^29^, thus, confirming our working hypothesis that aMIR46-48 is indeed silencing the *PUB46-48* genes.

**Fig. 2.**
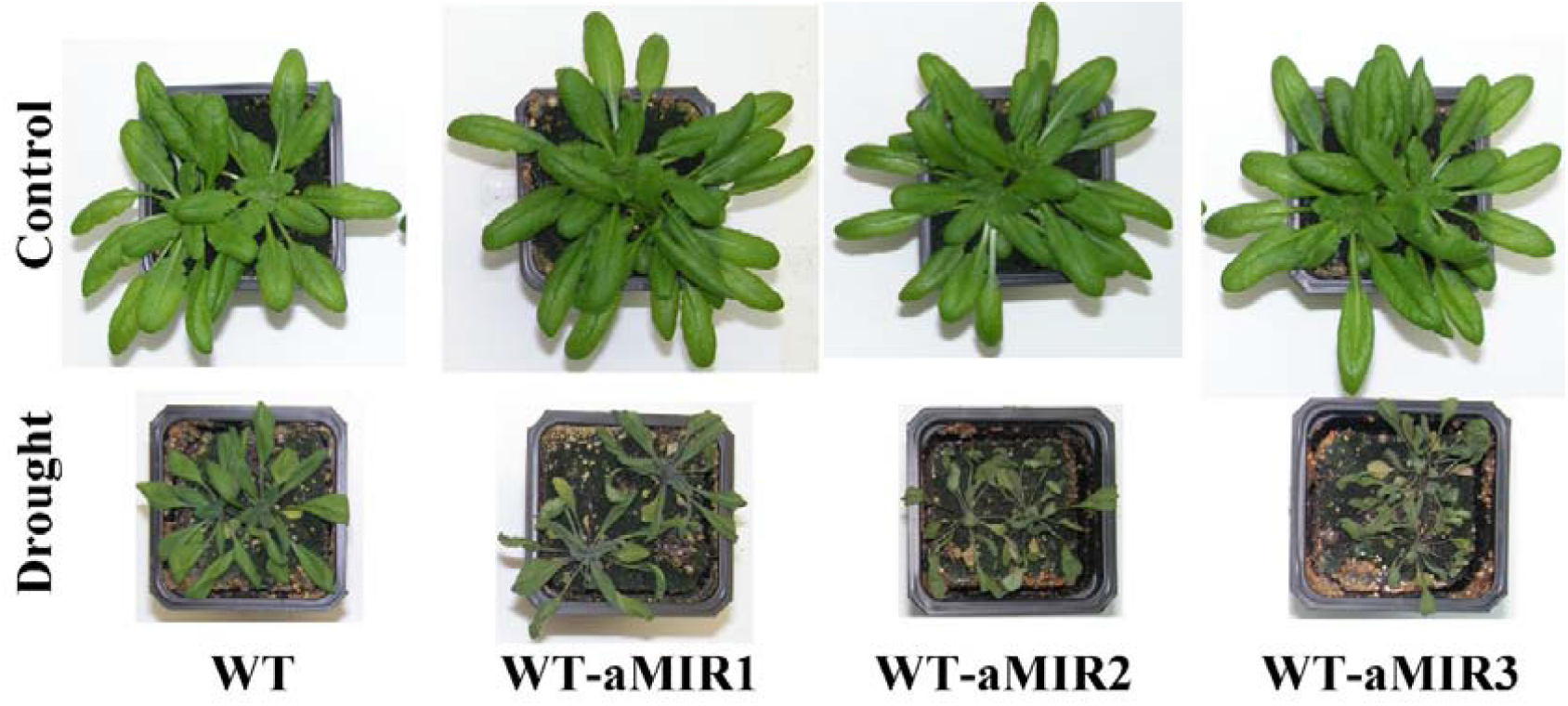
Water stress performance of WT-aMIR46-48 plants. WT plants and 3 lines of WT plants expressing aMIR46-48 were grown in pots for three weeks. Water was then withheld from drought treated plants, whereas control plants was watered. Plants were pictured 10 days after water withholding.

### *pub46* and *pub48* mutants overexpressing aMIR46-48

Here a direct comparison of single *pub46* or *pub48* mutant plants with or without aMIR46-48 expression enables us to test for an additive effect of severely downregulating the expression of all three genes, in essence creating a double/triple mutant plant. We determined sensitivity to drought stress, a phenotype observed in both *pub46* and *pub48* single gene mutants ^29^ and in WT plants transformed with aMIR46-48 (Fig. 2). Under optimal conditions there was no difference in the phenotypes of all the plants (Fig. S2). However, under water withholding stress *pub46* and *pub48* mutant plants expressing aMIR46-48 were far more drought sensitive than the parental single mutant lines or the WT-aMIR46-48 plants (Fig. 3). Drought stressed plants were rewatered and recovery was followed for the next 10 days. WT plants recovered best, followed by WT-aMIR46-48 plants. The *pub46* and *pub48* mutants struggled to recover whereas the *pub46* and *pub48* mutants that expressed aMIR46-48 did not recover from this stress at all (Fig. 4).

**Fig. 3.**
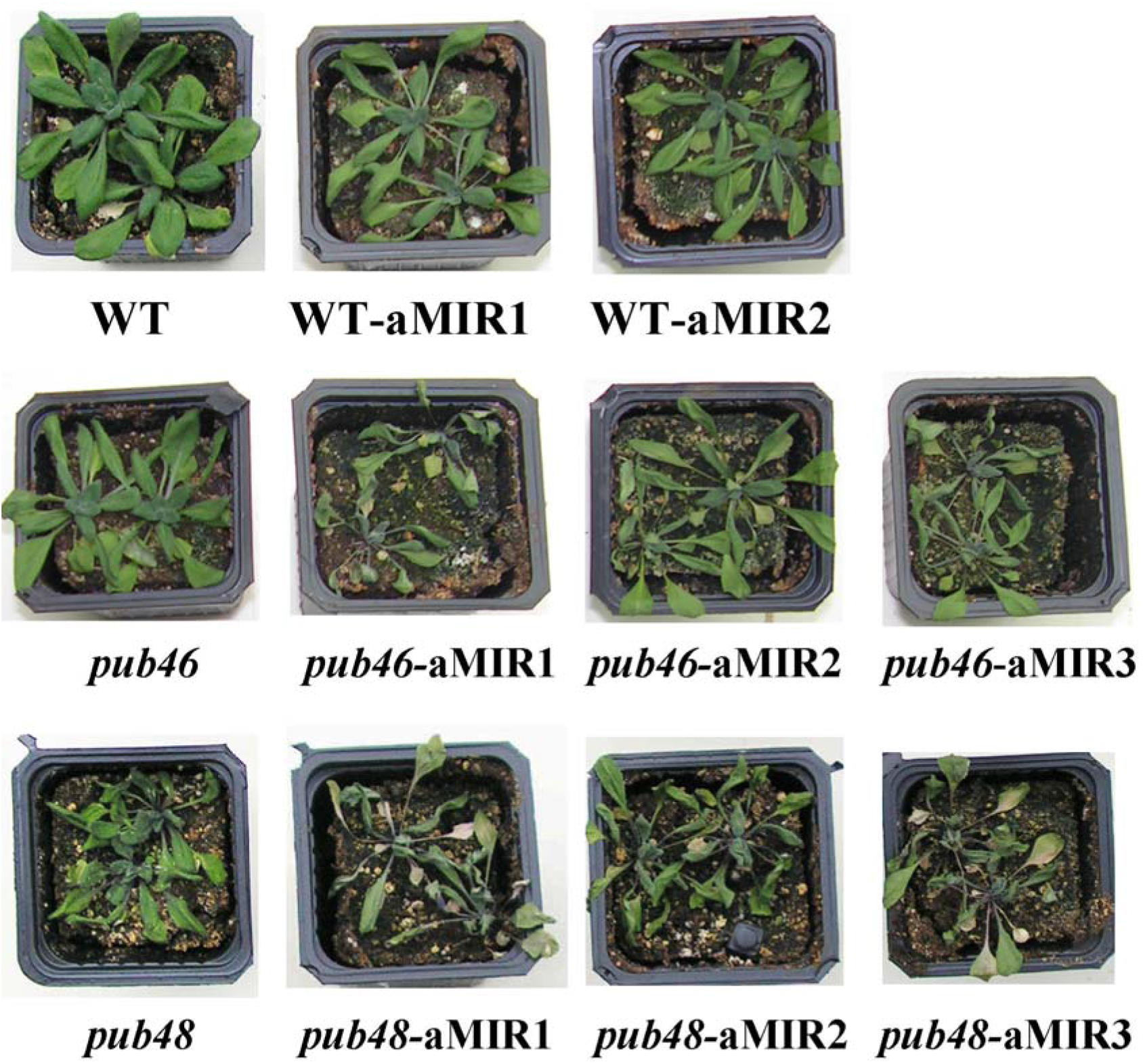
Phenotype of *pub46* and *pub48* mutants expressing aMIR46-48. Plants of the indicated genotype were grown in pots for 3 weeks. Water was then withheld for eight days. Control plants are shown in Fig. S2.

**Fig. 4.**
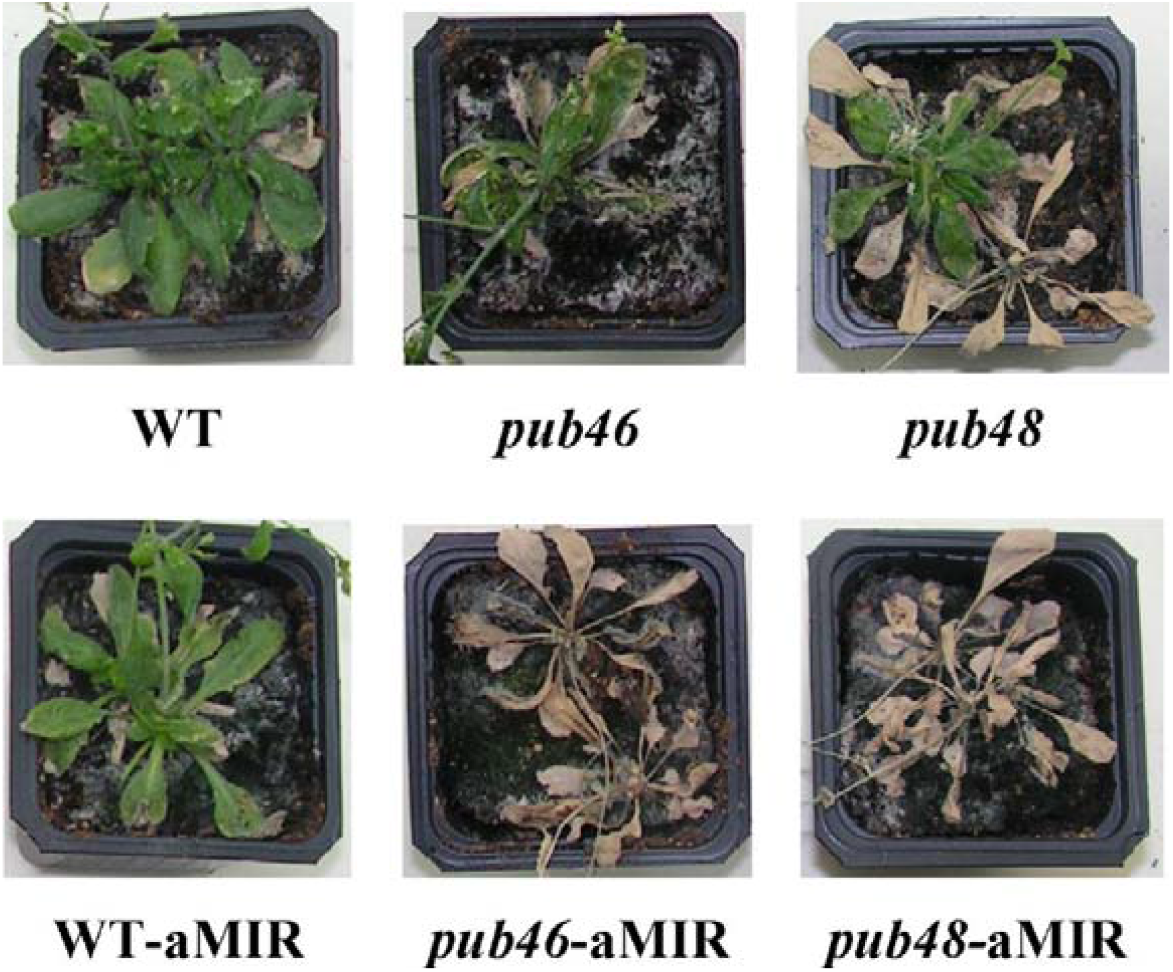
Plant recovery from drought stress. Plants treated as described in Fig. 3 were re-irrigated, and pictures were taken ten days later.

### Germination in the presence of ABA

The plant hormone abscisic acid (ABA) is the central hormone in drought and germination ^36^. We previously showed that germination in the presence of 1μM ABA of WT and single *pub46, pub47* or *pub48* mutants seeds was inhibited equally ^29^. This suggests that either the PUB46-PUB48 E3s are not involved in ABA signaling or their activity may be redundant. Here we addressed the role of functional redundancy by assaying the effects of ABA on germination of seeds of WT plants expressing aMIR46-48, in essence of the double/triple mutants. We found that these seeds were less sensitive to inhibition of germination and greening by ABA: 69-86% of seeds of the WT-aMIR46-48 line developed green seedlings 10 days after plating on medium containing 1 μM ABA as opposed to 45% of seeds of WT plants, suggesting that this activity of the PUB46-PUB48 E3s appears to be redundant (Fig. 5).

**Fig. 5.**
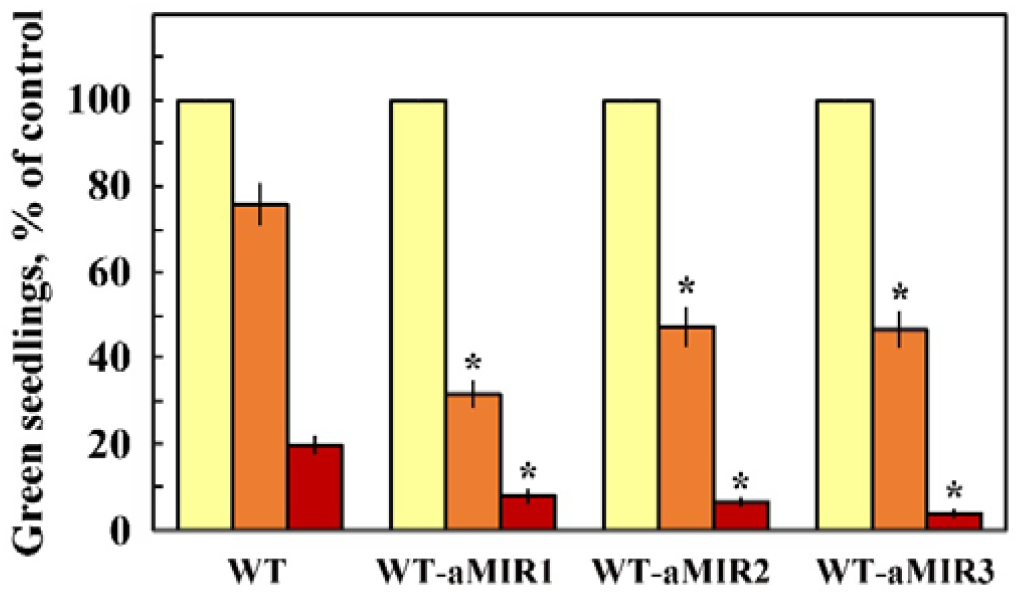
Effect of ABA on seedling germination. Surface sterilized cold treated seeds of WT and WT expressing aMIR46-48 were plated on agar media containing 0.5 X MS, 0.5% sucrose (yellow bars) supplemented with 1 μM (orange bars) or 1.5 μM (brown bars) ABA. Green seedlings were scored 7 days later. Data shown are average ± SE of 5 biological repeats with 40 seedlings for each treatment. Statistically significant changes from WT plants are marked with an asterisk.

### Germination in the presence of MV

Drought stress often results in oxidative stress ^37^. However, although each *pub46* or *pub48* single mutant is required for resilience to drought, we previously found that germination and seedling greening of *pub46* mutant seeds was hypersensitive to MV-induced oxidative stress whereas germination of *pub47* and *pub48* mutants resembled that of WT seedlings ^29^. We therefore tested the MV sensitivity of germination of WT-aMIR46-48 seeds. We found that WT-aMIR46-48 germination is hypersensitive to oxidative stress induced by MV (Fig. 6). For example, 28-43% of seeds of WT expressing aMIR46-48, developed green seedlings when germinated on 0.5 μM MV as opposed to 76% of the WT seeds.

**Fig. 6.**
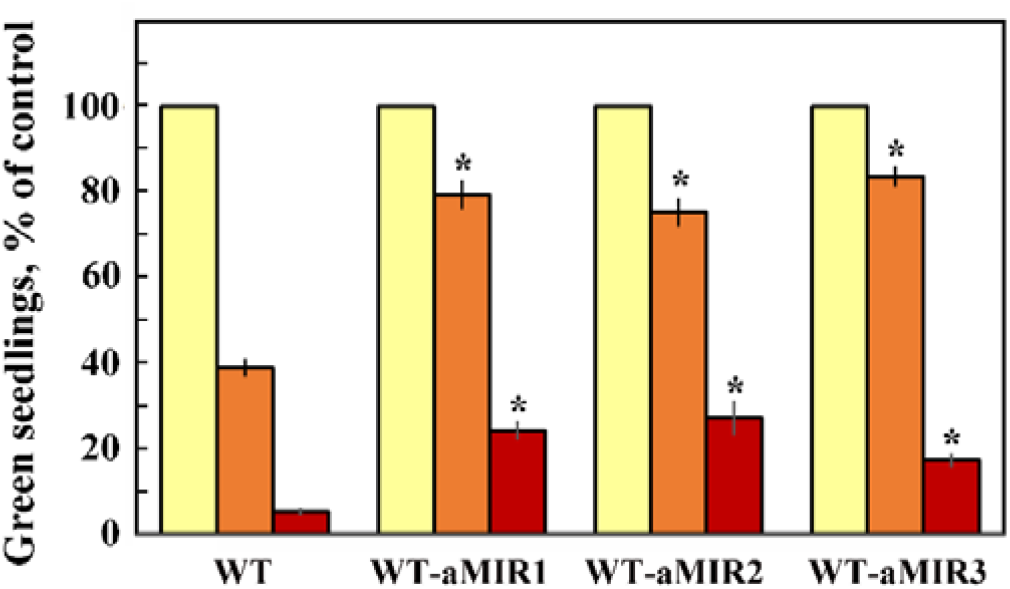
Effect of methyl viologen (MV). on seedling germination. Surface sterilized cold treated seeds of WT and WT expressing aMIR46-48 were plated on agar media containing 0.5 X MS, 0.5% sucrose (yellow bars) supplemented with 1 μM (orange bars) or 1.5 μM (brown bars) MV. Green seedlings were scored 7 days later. Data shown are average ± SE of 5 biological repeats with 40 seedlings for each treatment. Statistically significant changes from WT plants are marked with asterisk.

## Discussion

Our results indicate that RNA interference provides an efficient and accurate alternative means of severely reducing the activity of the entire *PUB46-48* gene family. This enabled us to address the question of the extent of redundancy in the function of these PUB E3s. The drought hypersensitivity of single *pub46* and *pub48* mutants ^29^ served as a reliable assay by which we now show that transformation of WT plants with aMIR46-48 recreates the drought hypersensitivity phenotype of each single mutant. This is a clear indication that the artificial sequence has indeed severely reduced the activity of the targeted *PUB46-48* genes. aMIR expression substantially reduces the expression of its target genes leaving some minor residual activity and a single aMIR targeted to multiple mRNAs may downregulate each target gene to a variable degree ^35^. Based on this result we then tested aMIR46-48 in single *pub46* and *pub48* mutants to differentiate between redundant and nonredundant functions of these E3s. The enhanced hypersensitivity of single *pub46* and *pub48* mutants to water withholding stress when they expressed aMIR46-48 indicates that each E3 performs unique function(s). Redundancy could indicate that each E3 participates in a single pathway whereas the nonredundant functions suggest that each E3 targets additional unique substrates for proteasomal degradation. Alternatively, the redundancy may be the result of partially overlapping temporal expression of these two PUBs.

Functional redundancy is a common feature of plant genomes with many genes belonging to gene families as a result of whole-genome duplication events ^31,32^. In the Arabidopsis *PUB* gene family there are various examples of redundancy in PUB E3 function: e.g., *pub4* mutants that have smaller rosette size show impaired stamen development resulting in lower fertility than that of WT plants ^38^ and increased proliferation of shoot and root meristems ^39,40^. In contrast, mutants of the highly homologous PUB2 E3 do not display any detectable phenotype ^41^. However, rosette size and stamen development of double *pub2 pub4* mutants were more pronounced than of pub4 mutants, suggesting partial redundancy in the function of PUB2 and PUB4 ^41^. Arabidopsis PUB12 and PUB13 ubiquitylate the brassinosteroid receptor BRI1 and the endosomal pool of BRI1 was reduced in *pub12 pub13* double mutants implying redundancy in their function ^42^. In addition, the Arabidopsis PUB22 and PUB23 E3s showed redundancy in modulating the degradation of the ABA receptor PYL9 ^43^. Furthermore, single mutants of homologous Arabidopsis *PUB25* and *PUB26* have weak freezing sensitivity compared to WT whereas *pub25 pub26* double mutants are hypersensitive to freezing, suggesting redundant function for these genes in ubiquitylation of the cold signaling negative regulator MYB15 ^44^. Arabidopsis PUB40 reduces the cellular concentrations of BRASSINAZOLE RESISTANT1 (BZR1) and BZR1 levels and the enhanced root growth phenotype of *pub39 pub40 pub41* triple mutant could be complemented by overexpressing *PUB40*, suggesting that these genes are redundant ^45^.

ABA is the key hormone involved in the plant response to drought ^46^. ABA also modulates germination and seedling greening (reviewed by ^47^) via various genes that encode transcription factors ^48-51^, participate in RNA metabolism ^52-55^ and via interaction with the ABA coreceptor-protein phosphatases 2C (PP2C)^56-58^ and their inhibitors ^59^. Ubiquitylation of ABA receptors and PP2Cs, by E3s from the PUB, EING, and RBR families plays a central role in modulating ABA signaling, germination and response to stress (reviewed by ^60,61^).

Functional redundancy in degradation of proteins involved in ABA signaling may occur between non-homologous E3s. For example, AFP1, DWA1/DWA2, KEG and ABD1 are all involved in the ubiquitylation of the ABI5 transcription factor ^62-65^. Similarly, a number of different E3s ubiquitylate the ABA receptors and co-receptors (reviewed by ^60^). Partial redundancy is also found in the signaling pathways of other plant hormones ^45,66^. In our experiments WT plants expressing aMIR46-48 were less sensitive to ABA inhibition of germination and seedling greening than the parental WT plants (Fig. 2). Interestingly, ABA sensitivity of the single *pub46, pub47* or *pub48* mutants is similar to that of the WT plants ^29^, suggesting that the ABA response may be redundant in these single E3 mutants.

Additional PUBs regulate ABA signaling by ubiquitylating ABA receptors and coreceptors (reviewed by ^61^: *pub 22 pub23* double mutants display enhanced tolerance to drought and target the ABA receptor PYL9 ^67^. In contrast, *pub12 pub13* double mutants are ABA-insensitive: PUB12 and PUB13 regulate ABA signaling by mediating the stability of the ABA co-receptor ABI1 ^68^. Interestingly, both PUB gene pairs display functional redundancy. Furthermore, PUB18 and PUB19 also showed redundancy in ABA inhibition of germination, as double *pub18 pub19* mutants were less sensitive compared to WT, whereas single mutants had similar ABA sensitivity to that of WT seeds ^69^. Our results suggest that PUB46 and PUB48 affect ABA signaling, the target is yet to be determined. If PUB46 and PUB48 affect ABA perception, it is likely that they target one or more of the ABA coreceptors, given that WT-aMIR46-48 germination displayed reduced ABA sensitivity, resembling that of the *pub12 pub13* double mutant ^68^.

Our results suggest that whereas *PUB46* and *PUB48* possess gene-specific biological activities ^29^, they also share some functional redundancy, *i*.*e*., their redundancy is partial. The extent of redundancy differs for the different activities/stresses. For example, redundancy can be observed in a more pronounced water stress hypersensitivity of a single gene mutant (Figs. 3 and 4), or in display of a phenotype such as germination in the presence of ABA observed solely when the activity of all the genes is reduced (Fig. 5). The partial functional redundancy of PUB46 and PUB48 may result from the protein substrate preferences of these E3s and their expression pattern in different plant tissues under various conditions. PUB46, PUB47 and PUB48 each contain 3 ARM motifs involved in protein-protein interaction, in particular with their ubiquitylation substrates. Each of the corresponding ARM motifs shares high sequence identity (60-71%) ^29^, suggesting that these ARM motifs are both close enough to bind the same proteins substrates, resulting in functional redundancy, as well as having differences that would enable them to target additional specific targets. Similarly, promoter activity of these 3 genes shows both overlapping and gene specific expression patterns ^29^, suggesting that gene expression patterns of paralogous genes may also contribute to the functional redundancy of their activity.

Classical models of gene duplication maintain that to survive through evolution, at least one of the paralogs must acquire a new function or else will be lost by deletion of nonfunctional genes at duplicate loci ^70-73^. The existence of truly redundant genes has been questioned ^74-76^, suggesting that all paralogs are likely to have some gene specific function. Our study joins other examples of partial redundancy of paralogous E3s and further comparative studies of the expression pattern of each paralog, their transcriptional, post transcriptional and post translational modifications, and their substrate preferences are needed to fully elucidate the complexity of the biological function of these E3s. Application of RNAi technology as we demonstrate here adds a new dimension to dissect the contribution of each E3 and to distinguish between overlapping redundant functions and those that are unique for each E3.

## Methods

### Plant material

*Arabidopsis thaliana* (Col). WT, *pub46* and *pub48* T-DNA insertion mutants were previously described ^29^.

### Construct for the expression of aMIR

Artificial MicroRNA (aMIR) was designed using the Web MicroRNA Designer (http://wmd.weigelworld.org/cgi-bin/webapp.cgi) ^35^. A synthetic DNA fragment with the appropriate 21 bp aMIR sequences was inserted into the MIR319a backbone ^35^ with flanking DNA sequences conferring *Xba*I and *Bam*HI sites for integration into the pCAMBIA 99-1 vector where expression is directed by the constitutive highly active 35S Cauliflower Mosaic Virus (35S CaMV) promoter.

### Plant Transformation

WT or the indicated T-DNA insertion mutant plants were transformed by the floral dip method ^77^ using *Agrobacterium* strain GV3101 harboring the respective plant transformation plasmid. Transgenic plants were selected on medium containing hygromycin, and homozygous plants resulting from independent transformation events by a single copy of the respective T-DNA were isolated as described ^77^.

### Plant growth and stress application

Plants were grown on agar plates with half-strength Murashige and Skoog (MS) nutrient solution ^78^ supplemented with 0.5% sucrose, or in planting mix in pots. Seed surface sterilization, imbibition, and growth conditions are detailed in ^29^. Where indicated, the growth medium was supplemented with the indicated concentrations of ABA or MV. Green seedlings were scored 7 days after plating. Each experiment was carried out in three biological repeats with treatment comprising ca 40 plants.

### Drought tolerance

For drought tolerance assays, seeds were planted in pots with equal amounts of potting mix. Plants were irrigated for three weeks, drought stress was applied by water withdrawal for the indicated time, and wilted plants were scored. Recovery was assayed by rewatering drought-treated plants and survival was scored ten days later.

## Acknowledgements

This work was supported by a grant No. 1352/17 from the Israel Science Foundation (ISF). DBZ is the incumbent of The Israel and Bernard Nichunsky Chair in Desert Agriculture, Ben-Gurion University of the Negev.

## Author contributions

GZV performed and analyzed the experiments and participated in writing the draft of the manuscript; DR contributed to experiments analysis and writing the manuscript; DBZ conceived the project idea, analyzed the experiments, and wrote the manuscript. All authors reviewed the manuscript.

## Ethics declarations

The authors declare no competing interests.

## Notes

### Competing Interest Statement

The authors have declared no competing interest.

## References

1 Vierstra, R. D. The ubiquitin-26S proteasome system at the nexus of plant biology. Nat. Rev. Mol. Cell Biol. 10, 385–397 (2009).

2 Glickman, M. H. & Raveh, D. Proteasome plasticity. FEBS Lett. 579, 3214–322 (2005).

3 Chen, L. & Hellmann, H. Plant E3 ligases: flexible enzymes in a sessile world. Mol. Plant 6, 1388–1404 (2013).

4 Koegl, M. et al. A novel ubiquitination factor, E4, is involved in multiubiquitin chain assembly. Cell 96, 635–644 (1999).

5 Tu, D., Li, W., Ye, Y. & Brunger, A. T. Structure and function of the yeast U-box-containing ubiquitin ligase Ufd2p. Proc. Natl. Acad. Sci. USA 104, 15599–15606 (2007).

6 Aravind, L. & Koonin, E. V. The U box is a modified RING finger - a common domain in ubiquitination. Curr. Biol. 10, R132–R134 (2000).

7 Wiborg, J., O’shea, C. & Skriver, K. Biochemical function of typical and variant Arabidopsis thaliana U-box E3 ubiquitin-protein ligases. Biochem. J. 413, 447–457 (2008).

8 Trujillo, M. News from the PUB: plant U-box type E3 ubiquitin ligases. J. Exp. Bot. 69, 371–384 (2018).

9 Gagne, J. M. et al. Arabidopsis EIN3-binding F-box 1 and 2 form ubiquitin-protein ligases that repress ethylene action and promote growth by directing EIN3 degradation. Proc. Natl. Acad. Sci. USA 101, 6803–6808 (2004).

10 Stone, S. L. et al. Functional analysis of the RING-type ubiquitin ligase family of Arabidopsis. Plant Physiol. 137, 13–30 (2005).

11 Lu, X. et al. Genome-wide identification and expression analysis of PUB genes in cotton. BMC Genom. 21, 213 (2020).

12 Kim, D. Y., Lee, Y. J., Hong, M. J., Kim, J. H. & Seo, Y. W. Genome wide analysis of U-Box E3 ubiquitin ligases in wheat (Triticum aestivum L.). Int. J. Mol. Sci. 22, 2699 (2021).

13 Sharma, B. & Taganna, J. Genome-wide analysis of the U-box E3 ubiquitin ligase enzyme gene family in tomato. Sci. Rep. 10, 9581 (2020).

14 Hu, D. K. et al. Genome-wide distribution, expression and function analysis of the U-Box gene family in Brassica oleracea L. Genes 10, 1000 (2019).

15 Song J M. X.,, Yang H, Yue L, Song J, Mo B. The U-box family genes in Medicago truncatula: Key elements in response to salt, cold, and drought stresses. PLoS ONE 12, e0182402 (2017).

16 Yu, Y. H. et al. Genome-wide identification and analysis of the U-box family of E3 ligases in grapevine. Russ. J. Plant Physiol. 63, 835–848 (2016).

17 Zeng, L. R., Park, C. H., Venu, R. C., Gough, J. & Wang, G. L. Classification, expression pattern, and E3 ligase activity assay of rice U-box-containing proteins. Mol. Plant.1, 800–815 (2008).

18 Tang, X. et al. Genome-wide identification of U-box genes and protein ubiquitination under PEG-induced drought stress in potato. Physiol. Plantar. in press (2021) doi:https://doi.org/10.1111/ppl.13475.

19 Wang, N. et al. Genome-wide identification of soybean U-Box E3 ubiquitin ligases and roles of GmPUB8 in negative regulation of drought stress response in Arabidopsis. Plant Cell Physiol. 57, 1189–1209 (2016).

20 Wang, C., Song, B., Dai, Y., Zhang, S. & Huang, X. Genome-wide identification and functional analysis of U-box E3 ubiquitin ligases gene family related to drought stress response in Chinese white pear (Pyrus bretschneideri). BMC Plant Biol. 21, 235 (2021).

21 Wang, K. L. et al. Genome-wide investigation and analysis of U-box ubiquitin-protein ligase gene family in apple: expression profiles during Penicillium expansum infection process. Physiol. Mol. Plant Pathol. 111, 101487 (2020).

22 Li, W. et al. Genome-wide and functional annotation of human E3 ubiquitin ligases Identifies MULAN, a mitochondrial E3 that regulates the organelle’s dynamics and signaling. PLOS ONE 3, e1487 (2008).

23 Andersen, P. et al. Structure and biochemical function of a prototypical Arabidopsis U-box domain. J. Biol. Chem. 279, 40053–40061 (2004).

24 Azevedo, C., Santos-Rosa, M. J. & Shirasu, K. The U-box protein family in plants. Trends Plant Sci. 6, 354–358 (2001).

25 Mudgil, Y., Shiu, S. H., Stone, S. L., Salt, J. N. & Goring, D. R. A large complement of the predicted Arabidopsis ARM repeat proteins are members of the U-box E3 ubiquitin ligase family. Plant Physiol. 134, 59–66 (2004).

26 Samuel, M. A., Salt, J. N., Shiu, S. H. & Goring, D. R. Multifunctional arm repeat domains in plants. Int. Rev. Cytol. 253, 1–26 (2006).

27 Zeng, L. R., Park, C. H., Venu, R. C., Gough, J. & Wang, G. L. Classification, expression pattern, and E3 ligase activity assay of rice U-box-containing proteins. Mol. Plant 1, 800–815 (2008).

28 Yee, D. & Goring, D. R. The diversity of plant U-box E3 ubiquitin ligases: from upstream activators to downstream target substrates. J. Exp. Bot. 60 (2009).

29 Adler, G. et al. The Arabidopsis paralogs, PUB46 and PUB48, encoding U-box E3 ubiquitin ligases, are essential for plant response to drought stress. BMC Plant Biology 17, 8 (2017).

30 Adler, G., Mishra, A. K., Maymon, T., Raveh, D. & Bar-Zvi, D. Overexpression of Arabidopsis ubiquitin ligase AtPUB46 enhances tolerance to drought and oxidative stress. Plant Sci. 276, 220–228 (2018).

31 Panchy, N., Lehti-Shiu, M. & Shiu, S. H. Evolution of gene duplication in plants. Plant Physiol. 171, 2294–2316 (2016).

32 Clark, J. W. & Donoghue, P. C. J. Whole-genome duplication and plant macroevolution. Trends Plant Sci. 23, 933–945 (2018).

33 Hung, Y. H. & Slotkin, R. K. The initiation of RNA interference (RNAi) in plants. Curr. Opin. Plant Biol. 61, 102014 (2021).

34 Voinnet, O. Origin, Biogenesis, and activity of plant microRNAs. Cell 136, 669–687 (2009).

35 Schwab, R., Ossowski, S., Riester, M., Warthmann, N. & Weigel, D. Highly specific gene silencing by artificial microRNAs in Arabidopsis. Plant Cell 18, 1121–1133 (2006).

36 Vishwakarma, K. et al. Abscisic acid signaling and abiotic stress tolerance in plants: A review on current knowledge and future prospects. Front. Plant Sci. 8, 12 (2017).

37 Scheibe, R. & Beck, E. Drought, Desiccation, and Oxidative Stress. in Plant Desiccation Tolerance (eds Lüttge, U., Beck, E. & Bartels, D.) pp. 209–231 Springer Berlin Heidelberg (2011).

38 Wang, H. et al. The Arabidopsis U–box/ARM repeat E3 ligase AtPUB4 influences growth and degeneration of tapetal cells, and its mutation leads to conditional male sterility. Plant J. 74, 511–523 (2013).

39 Kinoshita, A., Seo, M., Kamiya, Y. & Sawa, S. Mystery in genetics: PUB4 gives a clue to the complex mechanism of CLV signaling pathway in the shoot apical meristem. Plant Signal. Behav. 10, e1028707 (2015).

40 Kinoshita, A. et al. A plant U-box protein, PUB4, regulates asymmetric cell division and cell proliferation in the root meristem. Development 142, 444–453 (2015).

41 Wang, Y. P., Wu, Y. Y., Yu, B. Y., Yin, Z. & Xia, Y. J. EXTRA-LARGE G PROTEINs interact with E3 ligases PUB4 and PUB2 and function in cytokinin and developmental processes. Plant Physiol. 173, 1235–1246 (2017).

42 Zhou, J. et al. Regulation of Arabidopsis brassinosteroid receptor BRI1 endocytosis and degradation by plant U-box PUB12/PUB13-mediated ubiquitination. Proc. Natl. Acad. Sci. USA 115, E1906–E1915 (2018).

43 Zhao, J. et al. Arabidopsis E3 ubiquitin ligases PUB22 and PUB23 negatively regulate drought tolerance by targeting ABA receptor PYL9 for gegradation. Int. J. Mol. Sci. 18, 1841 (2017).

44 Wang, X. et al. PUB25 and PUB26 promote plant freezing tolerance by degrading the cold signaling negative regulator MYB15. Dev. Cell 51, 222–235 (2019).

45 Kim, E. J. et al. Plant U-Box40 mediates degradation of the brassinosteroid-responsive transcription factor BZR1 in Arabidopsis roots. Plant Cell 31, 791–808 (2019).

46 Zhu, J.-K. Abiotic stress signaling and responses in plants. Cell 167, 313–324 (2016).

47 Vishwakarma, K. et al. Abscisic acid signaling and abiotic stress tolerance in plants: a review on current knowledge and future prospects. Front. Plant Sci. 8, 161 (2017).

48 Giraudat, J. et al. Isolation of the Arabidopsis ABI3 gene by positional cloning. Plant Cell 4, 1251–1261 (1992).

49 Lopez-Molina, L. & Chua, N. H. A null mutation in a bZIP factor confers ABA-insensitivity in Arabidopsis thaliana. Plant Cell Physiol. 41, 541–547 (2000).

50 Finkelstein, R. R., Wang, M. L., Lynch, T. J., Rao, S. & Goodman, H. M. The Arabidopsis abscisic acid response locus ABI4 encodes an APETALA2 domain protein. Plant Cell 10, 1043–1054 (1998).

51 Wind, J. J., Peviani, A., Snel, B., Hanson, J. & Smeekens, S. C. ABI4: versatile activator and repressor. Trends Plant Sci. 18, 125–132 (2013).

52 Hugouvieux, V., Kwak, J. M. & Schroeder, J. I. An mRNA cap binding protein, ABH1, modulates early abscisic acid signal transduction in Arabidopsis. Cell 106, 477–487 (2001).

53 Lu, C. & Fedoroff, N. A mutation in the arabidopsis HYL1 gene encoding a dsRNA binding protein affects responses to abscisic acid, auxin, and cytokinin. Plant Cell 12, 2351–2365 (2000).

54 Nishimura, N. et al. Analysis of ABA Hypersensitive germination2 revealed the pivotal functions of PARN in stress response in Arabidopsis. Plant J. 44, 972–984 (2005).

55 Xiong, L. M. et al. Modulation of abscisic acid signal transduction and biosynthesis by an Sm-like protein in Arabidopsis. Dev. Cell 1, 771–781 (2001).

56 Leung, J., Merlot, S. & Giraudat, J. The Arabidopsis ABSCISIC ACID-INSENSITIVE2 (ABI2) and ABI1 genes encode homologous protein phosphatases 2C involved in abscisic acid signal transduction. Plant Cell 9, 759–771 (1997).

57 Meyer, K., Leube, M. P. & Grill, E. A protein phosphatase 2C involved in ABA signal transduction in Arabidopsis thaliana. Science 264, 1452–1455 (1994).

58 Nishimura, N. et al. ABA-Hypersensitive Germination1 encodes a protein phosphatase 2C, an essential component of abscisic acid signaling in Arabidopsis seed. Plant J. 50, 935–949 (2007).

59 Tischer, S. V. et al. Combinatorial interaction network of abscisic acid receptors and coreceptors from Arabidopsis thaliana. Proc. Natl. Acad. Sci. USA 114 (2017).

60 Ali, A., Pardo, J. M. & Yun, D.-J. Desensitization of ABA-signaling: the swing from activation to degradation. Front. Plant Sci. 11, 379 (2020).

61 Coego, A. et al. Ubiquitylation of ABA receptors and protein phosphatase 2C coreceptors to modulate ABA signaling and stress response. Int. J. Mol. Sci. 22, 7103 (2021).

62 Lopez-Molina, L., Mongrand, S., Kinoshita, N. & Chua, N. H. AFP is a novel negative regulator of ABA signaling that promotes ABI5 protein degradation. Gene Devel. 17, 410–418 (2003).

63 Lee, J.-H. et al. DWA1 and DWA2, Two Arabidopsis DWD protein components of CUL4-Based E3 ligases, act together as negative regulators in ABA signal transduction Plant Cell 22, 1716–1732 (2010).

64 Liu, H. & Stone, S. L. Abscisic acid increases Arabidopsis ABI5 transcription factor levels by promoting KEG E3 ligase self-ubiquitination and proteasomal degradation Plant Cell 22, 2630–2641 (2010).

65 Seo, K.-I. et al. ABD1 Is an Arabidopsis DCAF substrate receptor for CUL4-DDB1–based E3 ligases that acts as a negative regulator of abscisic acid signaling. Plant Cell 26, 695–711 (2014).

66 Hardtke, C. S. et al. Overlapping and non-redundant functions of the Arabidopsis auxin response factors MONOPTEROS and NONPHOTOTROPIC HYPOCOTYL 4. Development 131, 1089–1100 (2004).

67 Zhao, J. F. et al. Arabidopsis E3 ubiquitin ligases PUB22 and PUB23 negatively regulate drought tolerance by targeting ABA receptor PYL9 for degradation. Int. J. Mol. Sci. 18, 1841 (2017).

68 Kong, L. Y. et al. Degradation of the ABA co-receptor ABI1 by PUB12/13 U-box E3 ligases. Nature Commun. 6, 8630 (2015).

69 Bergler, J. & Hoth, S. Plant U-box armadillo repeat proteins AtPUB18 and AtPUB19 are involved in salt inhibition of germination in Arabidopsis. Plant Biol. 13, 725–730 (2011).

70 Ohno, S. Evolution by Gene Duplication. Berlin:Springer-Verlag (1970).

71 Nei, M. & Roychoudhury, A. K. Probability of fixation of nonfunctional genes at duplicate loci. Am. Nat. 107, 362–372 (1973).

72 Ohta, T. Evolution by gene duplication and compensatory advantageous mutations. Genetics 120, 841–847 (1988).

73 Walsh, J. B. How often do duplicated genes evolve new functionsã Genetics 139, 421–428 (1995).

74 Brookfield, J. Can genes be truly redundantã Curr. Bio. 2, 553–554 (1992).

75 Thomas, J. H. Thinking about genetic redundancy. Trends Genet. 9, 395–399 (1993).

76 Musso, G. et al. The extensive and condition-dependent nature of epistasis among whole-genome duplicates in yeast. Genome Res. 18, 1092–1099 (2008).

77 Clough, S. J. & Bent, A. F. Floral dip: a simplified method for Agrobacterium-mediated transformation of Arabidopsis thaliana. Plant J. 16, 735–743 (1998).

78 Murashige, T. & Skoog, F. A revised medium for rapid growth and bio assays with tobacco tissue cultures. Physiol. Plantar. 15, 473–497 (1962).

