## Supplementary material for "The Activity of the Stress Modulated Arabidopsis Ubiquitin Ligases PUB46 and PUB48 is Partially Redundant": Fig. S1, Fig. S2

TCTAGAGTCGACATACTCGCTGTTTTGAATTGATGTTTTAGGAATATATAT
GTAGAACCGATTAGAATACTGACTTTCACAGGTCGTGATATGATTCAATTA
GCTTCCGACTCATTCATCCAAATACCGAGTCGCCAAAATTCAAACTAGACT
CGTTAAATGAATGAATGATGCGGTAGACAAATTGGATCATTGATTCTCTTT
GATAGTCAGGATTCTAATCGCTTCTCTCTTTTGTATTCCAATTTTCTTGAT
TAATCTTTCCTGCACAAAAACATGCTTGGATCC

Supplementary Fig. 1. Nucleotide sequence of the aMIR46-48 construct.

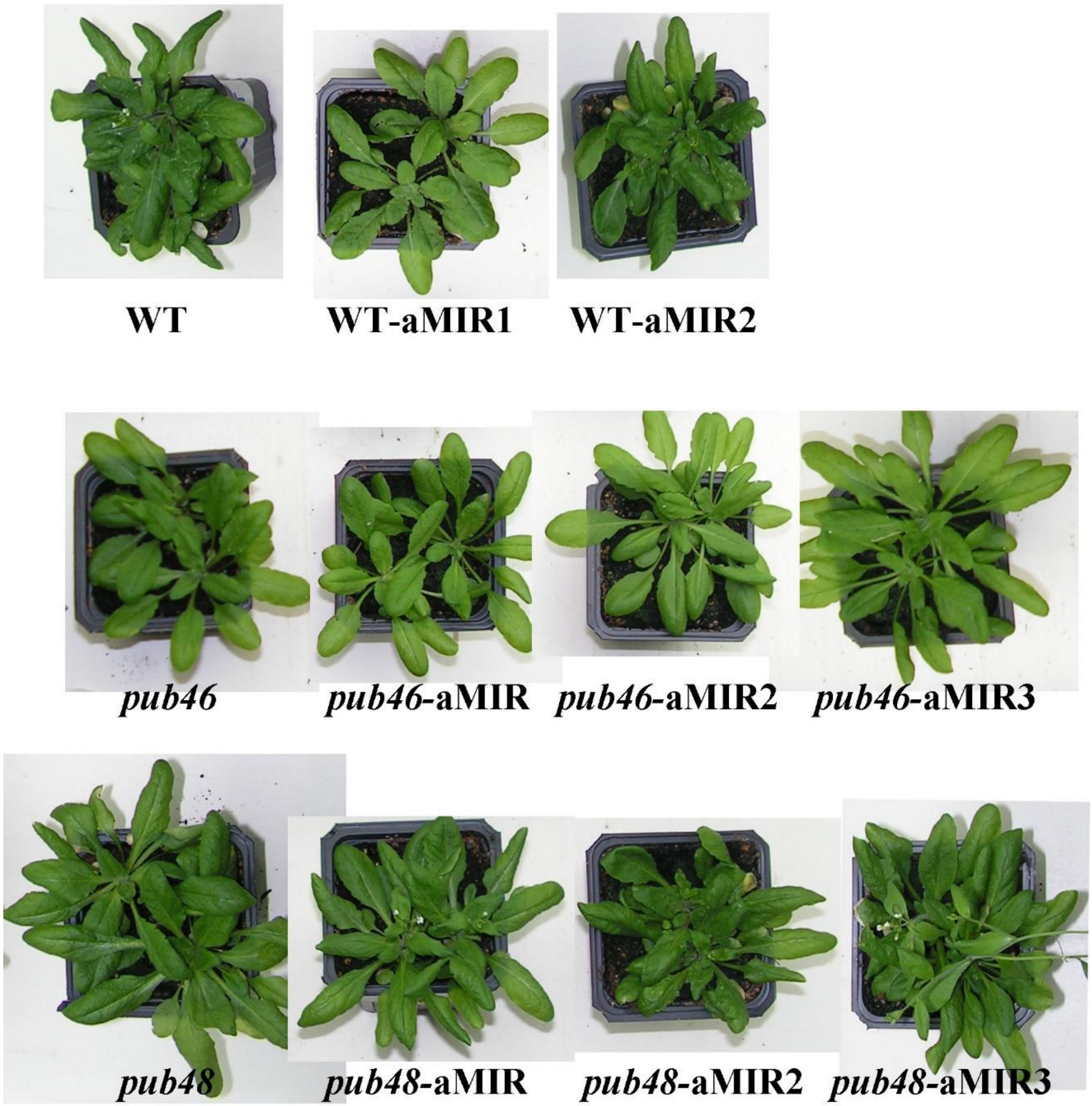


Supplementary Fig. 2. Irrigated control plants of the genotypes tested for drought stress shown in Fig. 3.
